# Tunneling nanotubes enable intercellular transfer in zebrafish embryos

**DOI:** 10.1101/2023.10.12.562005

**Authors:** Olga Korenkova, Shiyu Liu, Inès Prlesi, Anna Pepe, Shahad Albadri, Filippo Del Bene, Chiara Zurzolo

## Abstract

Tunneling nanotubes (TNTs) are thin intercellular connections facilitating the transport of diverse cargoes, ranging from ions to organelles. While TNT studies have predominantly been conducted in cell cultures, the existence of open-ended TNTs within live organisms remains unverified. Despite the observation of intercellular connections during embryonic development across various species, their functional role has not been confirmed. In this study, we performed mosaic labeling of gastrula cells in zebrafish embryos to demonstrate the coexistence of TNT-like structures alongside other cellular protrusions. These embryonic TNT-like connections exhibited similar morphology to TNTs described in cell culture, appeared to have similar formation mechanisms and could be induced by Eps8 overexpression and CK666 treatment. Most notably, to classify them as TNTs, we demonstrated their capability to transfer both soluble cargoes and organelles, which is a defining feature of open-ended TNTs. This study marks the first demonstration of functional TNTs in a living embryo.

## Introduction

Intercellular communication is a key feature of multicellular organisms. Along with soluble factors, cellular protrusions play an essential role in communication. Filopodia probe the environment, participate in cell-cell adhesion and chemoattractant sensing ^1^. Cytonemes are thin and long actin-based signaling protrusions that were first described in Drosophila wing imaginal disc ^2^, but were further shown to participate in signal transduction in other models, including tumors ^3^ and zebrafish embryo development ^4^. In addition to these mechanisms, more recently discovered structures – Tunneling nanotubes (TNTs), represent another type of contact-dependent communication.

TNTs are thin (below 1 µm) and long (up to 100 µm) actin-based membranous connections between cells, that allow for membrane and cytoplasmic continuity. Due to this feature, TNTs are able to transfer various material between connected cells: ions, mRNA, proteins, organelles, and, importantly, pathogenic agents, including misfolded proteins, viruses and bacteria ^5,6^. TNTs are suggested to participate in progression of cancer, Parkinson’s and Alzheimer’s diseases, in the spread of HIV and Sars-CoV2 ^7–11^. However, the majority of TNT studies are being performed in 2D cell culture, and there is still a limited number of studies that were performed in more complex 3D environment. Several studies have shown that TNT-like structures are present in resected tumors, tumor organoids, and developing embryos ^12–14^.

The challenge in identifying tunneling nanotubes (TNTs) within 3D models arises due to the absence of a specific TNT marker. Additionally, the prominent characteristic employed for TNT identification *in vitro*, namely the suspension of TNTs above the substrate, is not present within a 3D environment. In complex experimental settings, TNTs could be confused with other actin-based protrusions, including filopodia, cytonemes or cytokinetic bridges. The only reliable criteria to classify a structure as a TNT that is currently available, is to show that they are able to transfer cellular material. Consequently, structures observed in 3D models are often referred to as “TNT-like”, because their transfer functionality was not confirmed.

Another cellular structure that is able to transfer cellular materials is the intercellular bridge (IC). Differently from TNTs, intercellular bridges are cytoplasmic bridges that are formed by incomplete cytokinesis, keeping the cells interconnected after cell division. ICs are typically found in germline cells, where they enable communication via transfer of macromolecules, nutrients and organelles necessary for their further development ^15^. Interestingly, Caneparo and colleagues showed long and thin intercellular connections connecting cells in the zebrafish gastrula that were able to transfer cytoplasmic material^16^. The formation of these structures was associated with cell division and the authors suggested that they represent ICs that were formed by dividing cells during blastula stage and remained stable while cells were migrating apart in the gastrula.

Here we aimed to assess whether TNT-like structures (i.e., communication connections that do not arise from cell division) exist in a living embryo. We chose the early zebrafish embryo model for its optical transparency and rapid development that allows to follow *in vivo* processes in the living embryo in a non-invasive manner. Using a sparse cell labelling approach, we were able to discriminate and classify by morphology multiple types of protrusions formed by gastrula cells in the whole embryo, including functional TNT-like structures. We demonstrate that the latter not only share characteristics with TNTs observed in cell culture, but also have similar formation mechanisms, namely that they can be induced by overexpression of Eps8 (epidermal growth factor receptor kinase substrate 8) and by treatment with CK666. Most importantly, our results demonstrate that these connections have the ability to transfer intracellular material, such as cytoplasmic proteins and organelles. This study presents the first evidence of functional TNTs in a developing embryo in a whole organism context.

## Results

### Different types of protrusions co-exist in zebrafish gastrula

To visualize intercellular connections, zebrafish embryos were micro-injected with mRNA into one cell at the 16-cell stage, which resulted in mosaic expression of the construct at the gastrula stage (8 hpf) (Fig. 1A). Upon injection of membrane-labeling mRNA, protrusions with various morphologies could be observed within the same field of view (Fig. 1B). These included shorter, non-connecting protrusions, (arrow 1), longer protrusions, with a close-ended tip residing on the neighboring cell (arrow 2), connections with an interruption of membrane labelling often close to the middle of the connection (arrow 3), and uninterrupted connections, that appeared to link distant cells (arrows 4 and 5). The short protrusion (arrow 1) could be generally referred to as filopodia, while long closed-ended protrusions (arrow 2) were clearly distinguishable from filopodia and resembled classical cytonemes, that were previously observed at gastrula stage of zebrafish development ^4^ (Fig. 1C). On the other hand, the interruption in the connection’s labelling (arrow 3) hints towards the presence of midbody, thus this third type of connections could represent cytokinetic bridges, which could potentially span longer distances than canonical (1-5 µm) due to the migration of cells during gastrula stage (Fig. 1C). Finally, the uninterrupted connections between distant cells showed substantial variability in length and thickness and presented morphological characteristics of TNTs (Fig. 1C). Therefore, we quantified the connections as TNT-like structures only if they were above 5 µm in length ^17,18^ and appeared uninterrupted. We showed that at the gastrula stage, TNT-like structures connect about 35% of the labelled cells (Fig. 1D).

**Figure 1.**
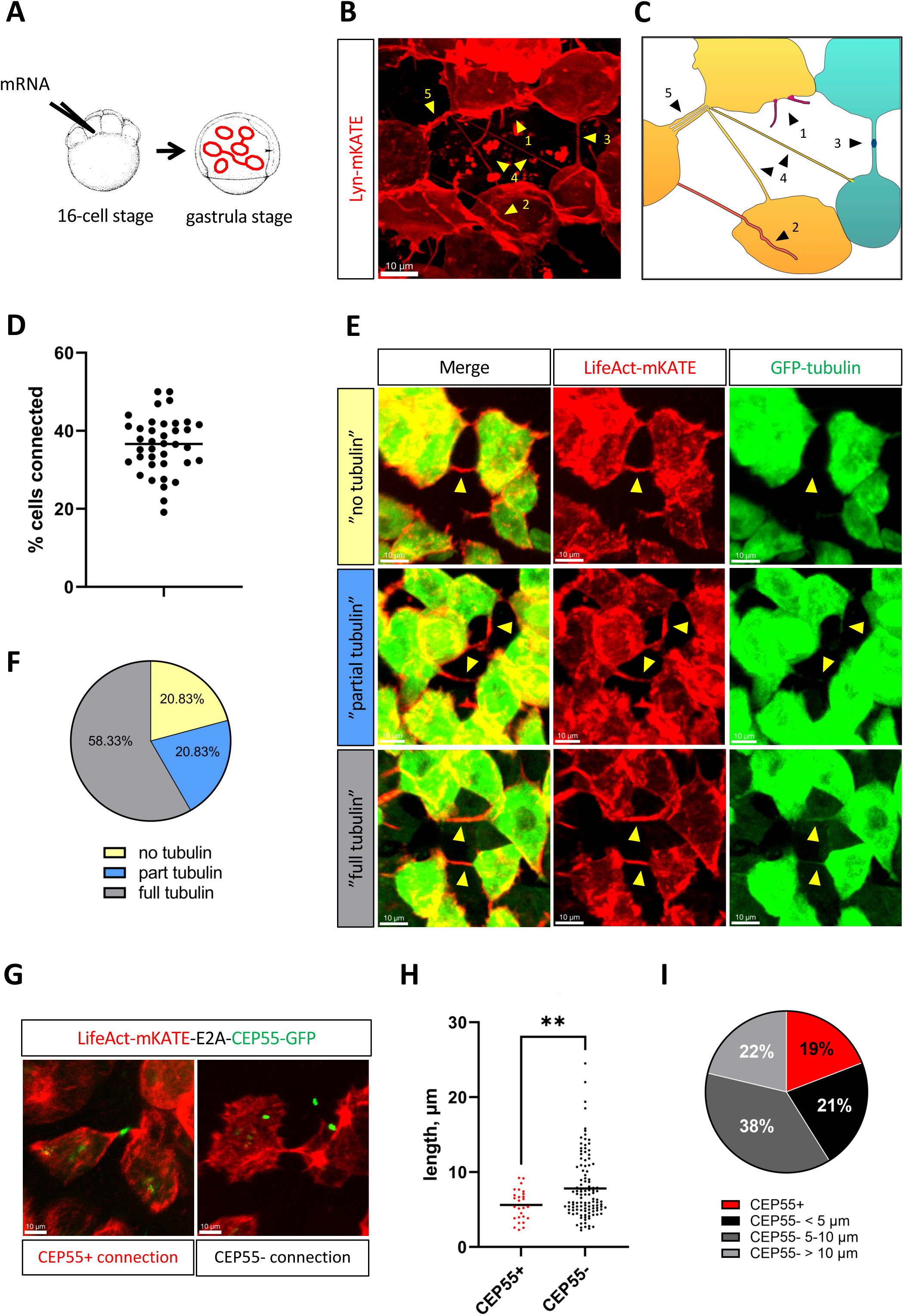
Characterization of intercellular connections in zebrafish gastrula. (A) Schematic representation of the labeling strategy to visualize intercellular connections: zebrafish embryo is injected into one of the central cells at 16-cell stage, incubated at 28°C until it reaches gastrula stage (7-9 hpf) and mounted for imaging. (B) Fluorescent photograph of different protrusions that can be observed at gastrula stage following membrane labeling by *lyn-mKATE* mRNA injection, pointed by yellow arrowheads: (1) short protrusions, (2) long closed-ended protrusions, (3) connections with interrupted membrane labeling, (4) thin TNT-like structures, (5) thick TNT-like structures. (C) Cartoon representing the image (B), suggesting the co-existence of: (1) filopodia, (2) cytonemes, (3) cytokinetic bridges, (4) TNT-like structures, (5) bundles of TNT-like structures. (D) Graph representing relative number of labelled cells, connected by TNT-like structures, each point representing separate embryo (n=38, median 36,6%). (E, F) Cytoskeletal composition of TNT-like structures. Embryos were injected as described in (A), using *lifeAct-mKATE-E2A-EGFP-tubulin* mRNA. All connections could be divided into 3 groups depending on the presence of tubulin. (E) – representative images, (F) – pie chart representing relative numbers of the 3 types of connections (n=72). (G, H) Presence of midbody in the intercellular connections. Embryos were injected as described in (A), using *lifeAct-mKATE-E2A-CEP55-EGFP* mRNA. (G) Representative images of CEP55-positive cytokinetic bridges and CEP55-negative TNT-like connections. (H) Graph representing difference in length distribution and average lengths between CEP55-positive (n=27, mean 5,6 μm) and CEP55-negative connections (n=114, mean 7,8 μm), p=0.0096, statistical analysis is t-test. (I) – pie chart representing relative numbers of CEP55-positive connections and CEP55-negative TNT-like connections having lengths below 5 μm, from 5 μm to 10 μm, and above 10μm (n=141). Scale bars are 10 μm.

Since TNTs *in vitro* can have different cytoskeletal composition depending on the cell type ^19^, in order to further characterize zebrafish TNT-like connections, we injected *lifeAct-mKATE-E2A-EGFP-tubulin* mRNA that labels actin and tubulin of the same cells (Fig. 1E). Quantification showed that the majority of TNT-like connections contained both actin and tubulin, while about 21% contained only actin (Fig. 1F).

### The majority of connections do not derive from cell division

To be able to differentiate the TNT-like connections we observed from cytokinetic bridges, we micro-injected *lifeAct-mKATE-E2A-CEP55-EGFP* mRNA that labels actin and the midbody marker CEP55 in the same cells (Fig. 1G). CEP55 was previously shown to label cytokinetic bridges in the zebrafish embryo throughout their lifespan, from the beginning of bridge formation until the midbody remnant release ^20^. In our timelapse recordings CEP55 was present from cytokinetic bridge formation until abscission, while CEP55-positive midbody remnants could also be observed (data not shown). We show that 19% of all identified connections were CEP55-positive and hence represented cytokinetic bridges (Fig. 1I), while the majority of connections were of a distinct nature. CEP55-positive cytokinetic bridges were significantly shorter than CEP55-negative connections, and their length never exceeded 10 µm (Fig. 1H), as expected for canonical cytokinetic bridges. Of interest, a portion of CEP55-negative connections had the length above 10 µm, and they could reach up to 30 µm in length (Fig. 1H, I).

A substantial amount of CEP55-negative connections had the length below 5 µm (Fig. 1I), which could correspond to filopodia that were reported to fall in this length range ^17,18^; thus, for further analysis of TNT-like structures we only included connections with a length above 5 µm.

Interestingly, in rare embryos we could observe unexpectedly long and stable cytokinetic bridges (Supplementary Fig.1). These structures exceeded 40 µm in length and remained stable for at least 30 minutes of the timelapse recording (Movie 1). While embryos containing these structures were rarely observed (around ten embryos from hundreds of embryos imaged in total), within these embryos several bridges could be seen in the same field of view. This observation suggests that these particular embryos experienced dysregulation during development, and the observed structures could possibly be DNA bridges formed as consequence of failed abscission ^21^. Overall, our approach allowed the discrimination of TNT-like structures from other intercellular connections in the embryo.

### Connections can be formed via two mechanisms similarly to TNTs in cell culture

Two mechanisms of TNT formation have been described *in vitro*: formation from filopodia-like structures that connect and fuse; and formation by cell dislodgement, when cells migrate apart and leave membrane tether behind ^5^. To be able to track formation of the TNT-like connections in the living embryo, we performed time-lapse experiments at 8 hpf. We were able to show that connections in the embryo can be formed from two filopodia-like structures, similarly to TNTs *in vitro* (Fig. 2A, Movie 2). Connections appeared very dynamic (example on Fig 2A was fully formed within 4 minutes) and remained stable for at least 5 minutes after formation (Movie 2). Moreover, we observed that connections could also be formed by cell dislodgement. Interestingly, in some instances TNT-like structures formed between cells remained connected while one of them underwent cell division (Fig. 2B, Movie 3). This is highlighted in Fig. 2B and Movie 3, which show the top cell (white asterisk) that migrates away from two bottom cells and leaves a membrane tether behind (yellow arrowhead). At the same time, top cells divides and both daughter cells maintain the connections with bottom cells. A cytokinetic bridge of typical morphology could also be observed between the two daughter cells (white arrowhead). This data further confirms that TNT-like structures can coexist with cytokinetic bridges, which are formed by distinct mechanism and hence represent a different type of connections.

**Figure 2.**
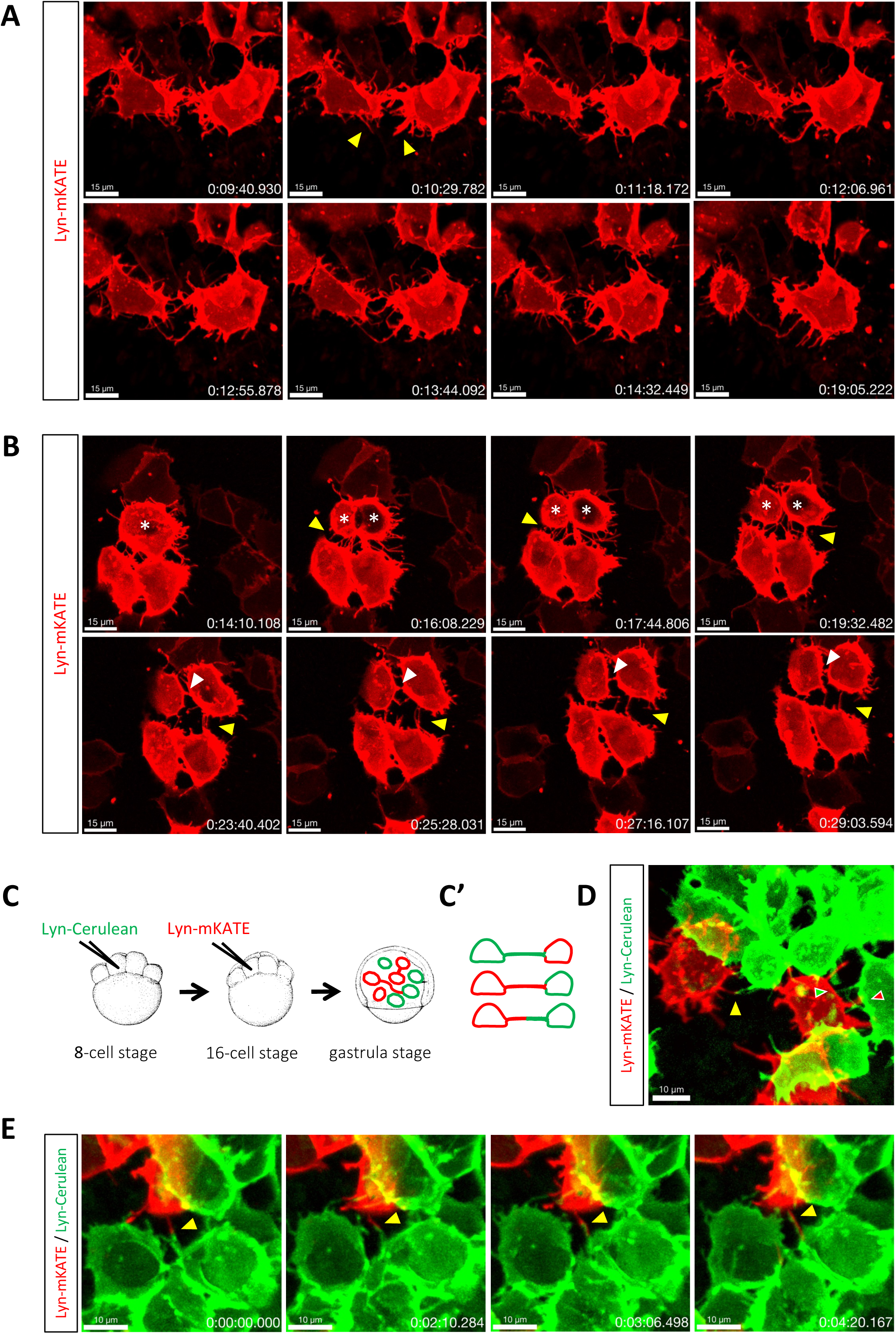
Formation mechanisms of TNT-like connections. (A) Time-lapse recording of the zebrafish embryo at 8hpf shows two neighboring cells extending filopodia (yellow arrowheads) that subsequently merge into single membranous connection that remains stable for at least 5 minutes. (B) Time-lapse recording of the zebrafish embryo at 8hpf shows three neighboring cells forming TNT-like connections by cell dislodgement (yellow arrowhead), while the top cell is dividing (white asterisks). Daughter cells maintain connections with lower cells for at least 15 minutes (yellow arrowhead), while cytokinetic bridge between daughter cells is also visible (white arrowhead). (C) Schematic representation of the strategy for labeling two cell populations: zebrafish embryo is injected into one of the central cells at 8-cell stage with *lyn-Cerulean* mRNA, followed by injection at 16-cell stage with *lyn-mKATE* mRNA, incubated at 28°C until it reaches gastrula stage (7-9 hpf) and mounted for imaging. (C’) – expected labeling outcomes: TNT-like structures formed by either one or both cell populations. (D) Representative image showing connections formed by either one (green and red arrowheads) or both (yellow arrowhead) cell populations. (E) Time-lapse recording of the zebrafish embryo at 8hpf shows two cells belonging to different cell populations, extending protrusions (yellow arrowhead) that elongate along each other. Scale bars are 15 μm for (A) and (B), 10 μm for (D) and (E), time is indicated as h:mm:ss.

As a complementary approach to confirm the formation of TNT-like structures through a mechanism distinct from cytokinesis, we conducted clonal labeling of zebrafish embryos. This involved injecting *lyn-Cerulean* mRNA into one of the cells at the 8-cell stage, followed by the injection of *lyn-mKATE* mRNA at the 16-cell stage (Fig. 2C). Using this method, two distinctly labeled cell populations were created and demonstrated that the connections could be labeled with a single color or both colors. Consequently, these connections could originate from either one or both of the connected cells (Fig. 2D). Interestingly, time-lapse recordings at 8 hpf showed that connections could be formed by elongation on top of each other, possibly serving each other as a support (Fig. 2E, Movie 4). This could lead to the formation of bundles of connections, which was previously shown to be one of the unique features of TNTs in cell culture ^22,23^.

### Eps8 and CK666 induce TNT-like structures in the embryo

Besides morphological criteria that allowed us to classify some of the connections observed in the embryo as TNT-like structures, we wanted to assess whether the latter were regulated by similar mechanisms as reported for TNTs *in vitro*. Our group has previously identified several ways to induce TNT formation in cell culture ^17,24^. One of the potent inducers of TNTs between cells in culture is Eps8, an actin binding protein with two distinct activities: actin-bundling and actin-capping. Upon overexpression of Eps8, the number of TNT-connected neuronal cells was increased, which was due to the bundling activity of the protein ^24^.

To overexpress Eps8 in the embryo, *eps8-HaloTag* mRNA was generated and micro-injected together with membrane-labeling *lyn-mKate* mRNA. Overexpression of the wildtype form of Eps8 (Eps8-WT) did not affect the number of TNT-connected cells but decreased average length of the connections (Fig. 3A, B, Supplementary Fig. 2A). This is probably due to dual activity of Eps8, namely, via its capping activity it could reduce the overall length of connections and hence the number of connections that we quantified following the length criteria. This hypothesis is supported by the fact that the number of shorter connections (5-10 µm) was increased (Fig. 3C).

**Figure 3.**
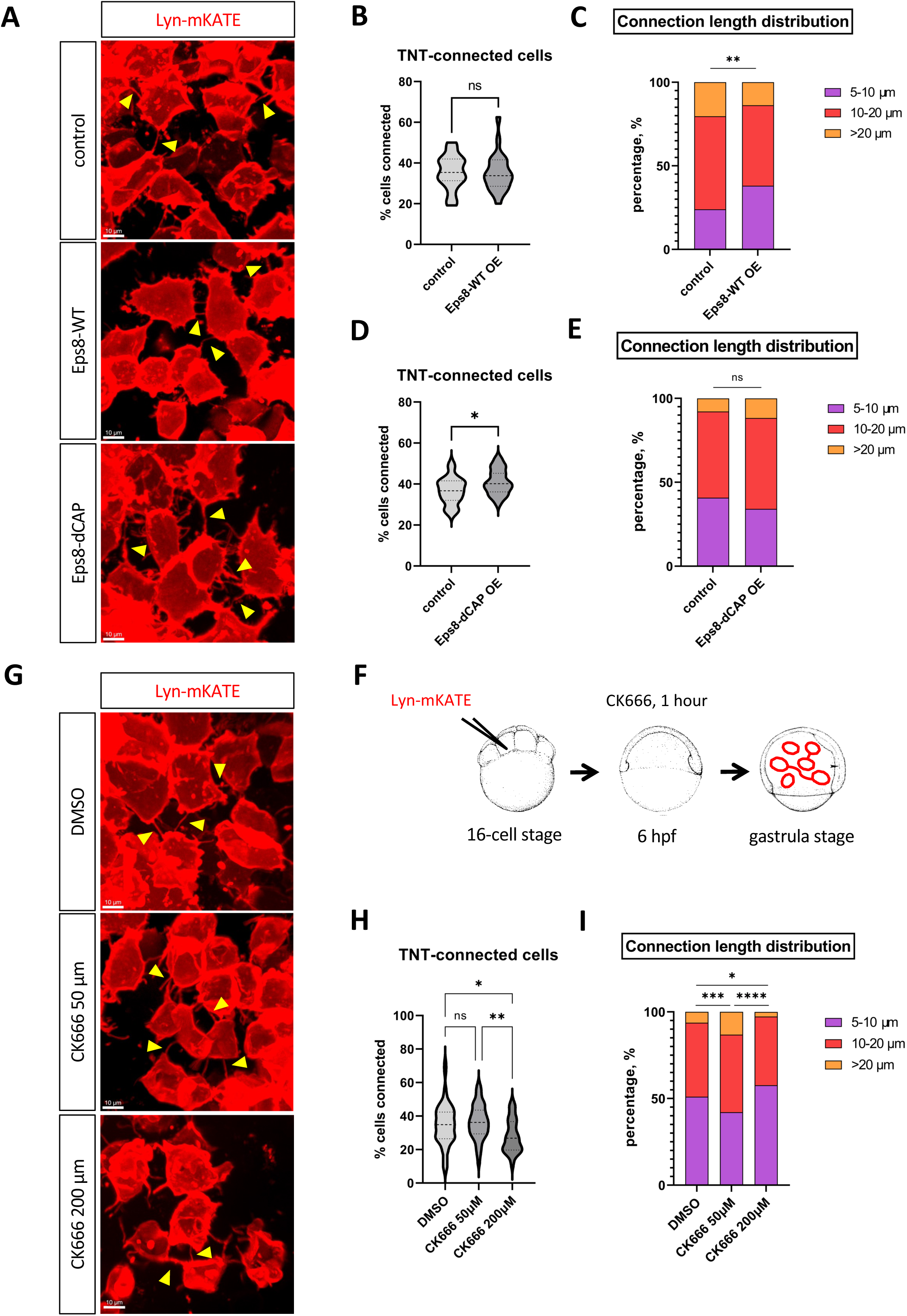
Effect of TNT-inducers on zebrafish intercellular connections. (A-E) Effect of overexpression of Eps8-WT (A-C) and its mutant Eps8-dCAP (A, D, E). (A) Representative images of the zebrafish embryo cells connected by TNT-like structures (yellow arrowheads), following Eps8-WT and Eps8-dCAP overexpression. Graphs (B) and (D) represent relative numbers of connected cells, graphs (C) and (E) represent length distributions of TNT-like connections. (B) control group: n=15, median 35.38%, Eps8-WT overexpression group: n=22, median 33.77%. p= 0.6521, analysed by Mann Whitney test. (C) control group: n=220, Eps8-WT overexpression group: n=247. p= 0.0030, analysed by chi-square test. (D) control group: n=23, median 36.71%, Eps8-dCAP overexpression group: n=28, median 40.19. p= 0.0436, analysed by Mann Whitney test. (E) control group: n=343, Eps8-dCAP overexpression group: n=489. p= 0.0516, analysed by chi-square test. (F-I) Effect of CK666 treatment of the embryo on TNT-like connections. (F) Schematic representation of experiment set-up: embryo was injected into one of the central cells at 16-cell stage, incubated at 28°C until it reaches 6hpf, treated with CK666 for 1 hour and mounted for imaging at gastrula stage (7-9 hpf). (G) Representative images of the zebrafish embryo cells connected by TNT-like structures (yellow arrowheads), following CK666 treatment. Graph (H) shows relative numbers of connected cells, graph (I) represents length distributions of TNT-like connections. (H) DMSO control group: n=57, median 34.88%, CK666 50 μM group: n=44, median 36.21%, CK666 200 μM group: n=24, median 26.81%. p-values are >0.9999 for DMSO vs. CK666 50μM, 0.0347 for DMSO vs. CK666 200μM and 0.0054 for CK666 50μM vs. CK666 200μM, analysed by Kruskal Wallis test. (I) DMSO control group: n= 541, CK666 50 μM group: n=328, CK666 200 μM group: n=298. p-values are 0.0008 for DMSO vs. CK666 50μM, 0.0297 for DMSO vs. CK666 200μM and <0.0001 for CK666 50μM vs. CK666 200μM, analysed by chi-square test.

Interestingly, when we overexpressed the mutant form of Eps8 lacking its capping activity (Eps8-dCAP), we observed an increase in the number of connected cells (Fig. 3A, D). Overexpression of Eps8-dCAP did not affect the length of connections (Fig. 3E, Supplementary Fig. 2B). These results are consistent with previous data where Eps8 was up-regulating TNTs in cell culture through its bundling activity ^24^.

To complement our characterization of TNT-like structures in the embryo in comparison with data obtained *in vitro*, we used a drug which differentially affected TNTs and other cell protrusions in cultured cells ^17,22^. Specifically, treatment of cells with the drug CK666, which inhibits Arp2/3 protein and hence branched actin network ^25^, was shown to induce TNT formation in neuronal cells while inhibiting attached filopodia and lamellipodia ^17,22^. To check whether CK666 would have a similar effect on the connections in the embryo, we incubated embryos with two different concentrations of the drug that were previously used in cell culture, for 1 hour prior to imaging (Fig. 3F). At 200 µM, CK666 appeared to be toxic to the embryos, that were either dying at early stages or, if survived, had observable tail twist at 48 hpf (Supplementary Fig. 3). This deleterious treatment resulted in the decrease of the number of connected cells and the average length of connections (Fig. 3G, H, I, Supplementary Fig. 2C). At 50 µM, CK666 was not altering the gross morphology of the embryo at the same developmental stage and the overall survival. However, at this dose the treatment led to an increase of the average length of connections as well as the number of longer connections (above 10 µm), while the total number of connected cells was not affected (Fig. 3G, H, I, Supplementary Fig. 2C). This data is consistent with the role of Arp2/3 as a TNT inhibitor in cell culture ^17^. However, as Arp2/3 participates in the cytokinetic bridge abscission ^26^, we next questioned whether the length of cytokinetic bridges would also be affected by CK666 treatment. Incubation of *LifeAct-mKATE-E2A-CEP55-EGFP* –injected embryos with 50 µM CK666 resulted in a significant increase in the average length of cytokinetic bridges (from an average of 4.7 µm in DMSO-treated embryos to an average of 5.7 µm in CK666-treated embryos). Nevertheless, CEP55-positive connections still did not exceed 10 µm in length (Supplementary Fig. 2D). These observations indicate that Arp2/3 inhibition had a prominent effect on TNT-like connections in zebrafish similar to what has been shown in *in vitro* cell culture studies ^17,22^.

### TNT-like structures are able to transfer material

The most important criteria to define TNTs is to assess their transfer function. Thus, we next investigated whether the characterized TNT-like connections *in vivo* could transfer material and hence represent functional TNTs. To this purpose we set up several approaches. First, to be able to observe the transfer of cytoplasmic proteins, we made use of the Dendra2 photoconvertible protein. The molecular weight of Dendra2 protein (26 kDa) gives a possibility to rule out gap junction –mediated transfer that has a cutoff of 1 kDa. Embryos were injected with *dendra2* mRNA along with *lyn-HaloTag* mRNA, which allowed us to visualize membranous connections (Fig. 4A, indicated by a yellow arrowhead) and subsequently photoconvert one of the connected cells (marked with a white asterisk) from green-Dendra2 to red-Dendra2. Time-lapse recording revealed that within 20 minutes, the connected cell (indicated by a white arrowhead) received red labeling from the photoconverted cell, while the fluorescent intensity of not connected neighboring cells (1, 2, 3) remained unchanged (Fig. 4A, B, Movie 5). Moreover, linear regression curve fitting showed that the photoconverted donor cell was losing red fluorescent signal with similar speed as the acceptor cell was receiving the red signal (slope = –0,46; R^2^ = 0,86 for donor cell, slope = 0,49; R^2^ = 0,89 for acceptor cell) (Fig. 4B). This first approach therefore reveals active transfer of Dendra2 photoconverted protein between the two cells through the Lyn-HaloTag –labeled open-ended TNT connection.

**Figure 4.**
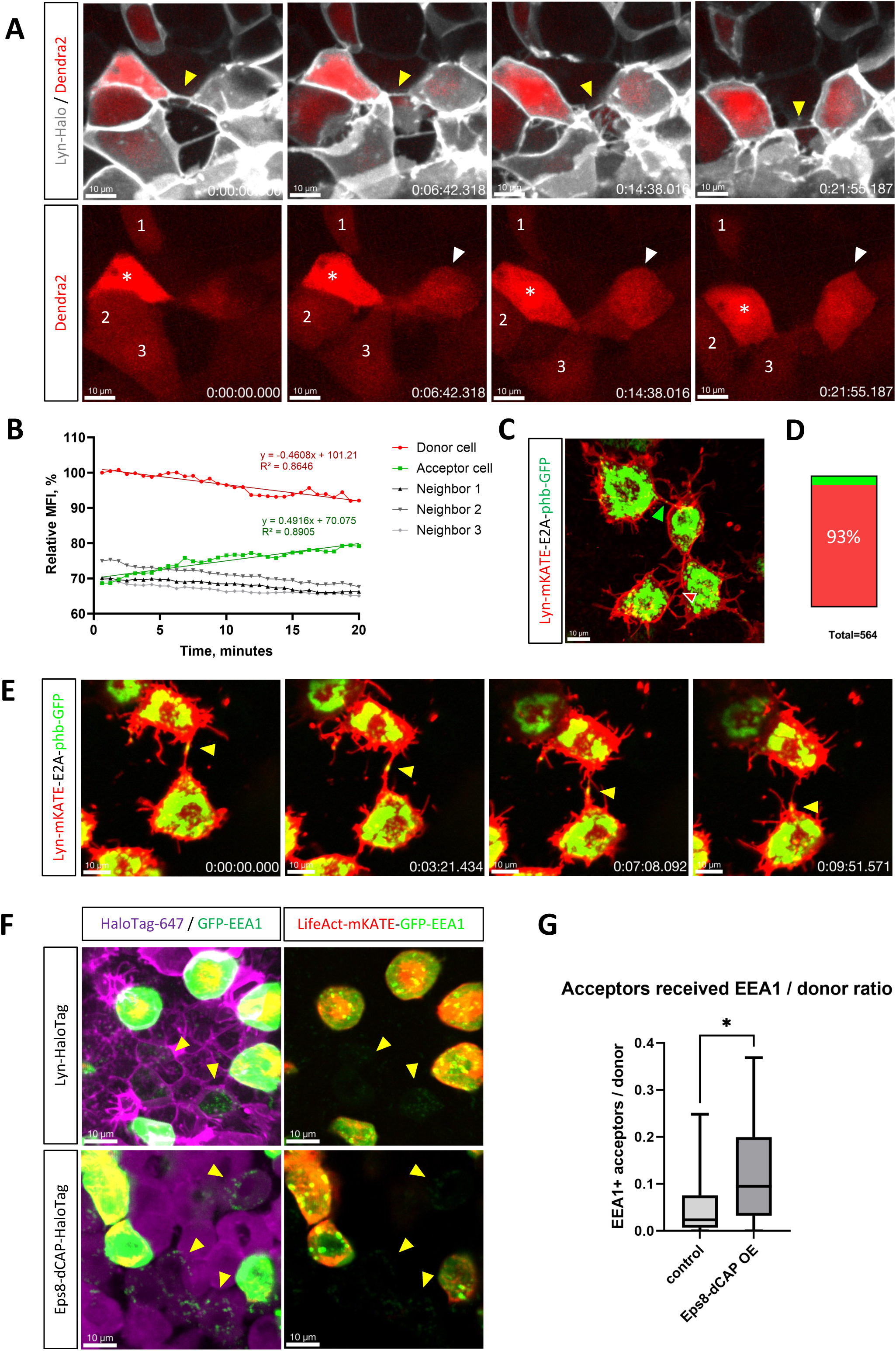
TNT-like structures are able to transfer material. (A) Time-lapse recording of the zebrafish embryo at 8hpf shows two cells connected by a membranous TNT-like connection (yellow arrowhead). Cytoplasmic Dendra2 was photoconverted to red in the cell on the left (white asterisk), recording shows that cell on the right (white arrowhead) receives red labeling within 20 minutes, while the fluorescence intensity of other neighboring cells (1, 2, 3) is not changed. (B) Relative mean fluorescent intensity (MFI) quantification for timelapse (A). Linear regression fitting for donor and acceptor cells performed in GraphPad Prism software, equations and goodness of fit (R^2^) visualized on the graph. (C) Representative image of the zebrafish embryo cells connected by TNT-like structures, where some connections contain mitochondria (green arrowhead). Red arrowhead points at TNT-like structure quantified as not containing mitochondria. (D) Quantification of the relative number of TNT-like connections containing mitochondria (red part – no mitochondria, green part – containing mitochondria), n=564. (E) Time-lapse recording of the zebrafish embryo at 8hpf shows two cells connected by a membranous TNT-like connection which enable the transfer of mitochondria (yellow arrowhead). (F) Representative images showing magenta Lyn-HaloTag 647 or Eps8-dCAP-HaloTag 647 acceptor cells containing green dots of EEA1-positive vesicles (yellow arrowheads). Green signal on the left panels was enhanced compared to the right panels, in order to better visualize vesicles in acceptor cells. (G) Quantification of the relative amount of acceptor cells (normalized to the amount of donor cells) that received EEA1-positive vesicles. Control group: n=17, median 0.02344, Eps8-dCAP overexpression group: n=18, median 0.09467. p= 0.0184, analysed by Mann Whitney test. Scale bars are 10μm, time is indicated as: h:mm:ss.

Next, to test whether TNT-like structures are able to transfer organelles, we injected *Lyn-mKATE-E2A-phb-EGFP* mRNA that would label membrane and mitochondria of the same cells (Fig. 4C) and therefore enable assessment of the ability of TNTs to transfer mitochondria. Quantification showed that approximately 7% of the TNT-like structures contained mitochondria, indicating the possibility of their transfer between cells (Fig. 4D). Time-lapse recording indeed confirmed that mitochondria could move along membranous TNT-like structures and be transferred between connected cells (Fig. 4E, Movie 6).

Finally, we investigated whether TNT-like structures could facilitate the transfer of vesicular structures. To track early endosomes, we labeled them using the Early Endosome Antigen 1 (EEA1) marker. For this experiment, we injected either *lyn-HaloTag* (control) or *eps8-dCAP-HaloTag* mRNA into the embryos at the 1-cell stage, followed by the injection of *lifeAct-mKATE-E2A-EGFP-EEA1* mRNA at the 16-cell stage. Subsequently, injected embryos were treated with the HaloTag 647 ligand prior to imaging. As result, two distinct cell populations were observed: donor cells with membrane (or cytoplasm), actin, and endosome labeling, and acceptor cells with only membrane (or cytoplasm) labeling (Fig. 4F). Timelapse experiments at 8 hpf showed that early endosomes can indeed be transferred via TNTs between labelled cells (Movie 7). Steady images of these injected embryos revealed LifeAct-mKate –negative, EGFP-EEA1-positive acceptor cells, indicating that they received early endosomes from LifeAct-mKate/EGFP-EEA1 –positive donor cells (Fig. 4F, yellow arrowheads). Furthermore, quantification showed that the relative number of acceptor cells that acquired early endosomes is significantly increased upon overexpression of Eps8-dCAP (Fig. 4G). Since overexpression of Eps8-dCAP was also increasing the number of TNT-connected cells (Fig. 3D), we conclude that the observed early endosome transfer is TNT-related.

## Discussion

TNT studies *in vivo* are very challenging due to the absence of specific markers, which makes it necessary to rely majorly on the functional characterization. The main morphological criterion to classify a connection as a TNT in cell culture studies is that TNT should be hovering above and not attach to the substrate. This positional criterion cannot be used in 3D models because the cells are embedded in extracellular matrix, which represents substrate of 2D models. Another complication of TNT visualization *in vivo* is that cells are very densely packed, which makes it impossible to observe intercellular structures if all of the cells are labelled. In our study we took advantage of the zebrafish embryo model which allows mosaic labeling of the embryonic cells following micro-injection of mRNA into one of the cells at 8-16 cell stage. We have chosen micro-injection into one of the central cells at 16-cell stage to achieve optimal labeling density.

Following micro-injections of different labeling mRNAs, we concluded that different types of protrusions can co-exist during zebrafish embryonic development. Analysis of the protrusions’ morphology suggested co-existence of filopodia, cytokinetic bridges, cytonemes and TNT-like structures within the same cell population of zebrafish gastrula cells. We then used additional criteria to the solely membrane labelling approach in order to clearly discriminate the different observed protrusions from TNT-like structures.

Gastrula cells are expected to form multiple filopodia as they are actively migrating during gastrulation. However, filopodia generally does not exceed 5 µm in length ^1,17^, so we used the criteria of length threshold to discriminate TNT-like connections from filopodia.

In parallel to active migration, gastrula cells are continuously dividing, so other frequently observed structures are cytokinetic bridges. Although conventional cytokinetic bridges should not exceed 5 µm in length, similarly to filopodia, there is a possibility that they get elongated when daughter cells migrate apart from each other, as previously hypothesized^16^. Therefore, cytokinetic bridges might be difficult to discriminate from TNT-like structures only based on length. Here, we have used the CEP55 midbody marker to distinguish TNT-like structures from cytokinetic bridges and have shown that the majority of intercellular connections (81%) did not derive from cell division. Based on our observations, CEP55-positive cytokinetic bridges never exceeded 10 µm in length, while TNT-like connections could span several tens of micrometers. However, out of hundreds of embryos imaged overall, in some embryos we were able to observe unexpectedly long and stable cytokinetic bridges (Supplementary Fig.1). Given that these structures could be observed between multiple daughter cells within the same embryo, we hypothesize that these particular embryos might have had alterations in development and the observed bridges could represent DNA bridges that are formed between daughter cells due to chromosome mis-segregation, and failed abscission ^21^.

Cytonemes, closed-ended structures that morphologically resemble TNTs ^2,27^, were shown to exist during zebrafish gastrulation, where they participate in Wnt signaling ^28^. In our study, we also observed cytoneme-like protrusions that appeared to be closed-ended. Consequently, we did not include them in the quantification of TNT-like protrusions, and only considered protrusions connecting two labeled cells and displaying continuous membrane labeling. However, solely relying on this morphological criterion, we cannot rule out the possibility that some structures counted as TNT-like might actually be cytonemes, as we lacked specific exclusion markers. In fact, similarly to TNTs, specific marker for cytonemes has not yet been identified.

Thus, we pursued our analysis to address whether the TNT-like connections, that in the embryo appeared morphologically distinct from filopodia, cytokinetic bridges and cytonemes, could represent actual TNTs that were well characterized in cell culture.

TNTs were initially described as actin-based protrusions that lack microtubules ^29^. However, further studies in other cell types showed that TNTs can also contain tubulin ^30,31^. We found that the majority of zebrafish TNT-like connections contained both actin and tubulin, while a proportion of connections contained only actin.

Next, we assessed the mechanism of TNT-like structures formation in the embryo and investigated whether they could be regulated by some of the factors identified for TNTs in cell culture. By timelapse microscopy, we found that both mechanisms previously described in cell culture, namely filopodia extension and cell dislodgement ^5,14^, lead to the formation of TNT-like structures between gastrula cells. Moreover, we could show that upon formation TNT-like connections tend to serve as a support for each other extension which might lead to the formation of bundles of connections. Similar observation has been reported in living cells in culture ^23^, while bundle organization has been described for TNTs in neuronal cells ^22^. Interestingly, we observed that TNT-like structures can form in parallel with cell division and cytokinetic bridge formation, which further supports the idea that distinct types of protrusions can coexist simultaneously within the same cells.

To assess whether intercellular connections that we observed in zebrafish embryos could be influenced by the effectors that were shown to induce TNT-formation in cell culture, we decided to investigate the role of Eps8, an actin-binding protein that is expressed during gastrulation ^32^ and was previously shown to induce TNT-formation in neuronal cells through its bundling activity ^24^. Eps8 displays two actin-regulating activities: capping actin barbed ends and cross-linking actin filaments ^33^. Overexpression of Eps8-WT, which possesses both activities, led to an increase in the number of shorter connections, probably because under WT conditions its preferred activity is capping actin filaments. In addition, by not considering structures less than 5 µm in length, we may have overlooked changes in the total amount of TNTs because the overall length of connections decreased. On the other hand, when Eps8-dCAP mutant (lacking capping activity) was overexpressed, the number of cells connected by TNT-like structures was increased. This is consistent with studies on the effect of Eps8 on TNTs in cell culture and might be due to an increase of the TNT stability and/or to an increase of TNT formation induction as previously reported ^17,24^.

As an alternative approach to affect TNT formation in the whole embryo, we used CK666, a drug that inhibits Arp2/3 branched actin network ^25^ and robustly induces TNTs in culture^17^. Interestingly, we found that it significantly increased the proportion of longer TNT-like connections. This is in line with cell culture data where CK666 releases the pool of actin from branched actin network and hereby enables formation of longer connections ^17,22^. Arp2/3 has also been demonstrated to participate in cytokinetic bridge abscission ^26^. Therefore, inhibiting Arp2/3 could potentially influence the stability of cytokinetic bridges. We hypothesized that if cytokinetic bridges persist longer, their length might increase due to continuous cell migration during gastrulation. As expected, we observed that CK666 treatment increased the average length of CEP55-positive cytokinetic bridges, although they still did not exceed 10 µm in length. Interestingly, CK666 only affected TNT-like connections that were above 10 µm in length, indicating that elongated cytokinetic bridges did not interfere with the results obtained for TNT-like connections.

Taken together, our data suggests that TNT-like intercellular connections in the zebrafish embryo share multiple characteristics with TNTs observed in cell culture, namely: i) a similar morphology, ii) similar formation mechanisms, and iii) similar response to known TNT inducers. However, the only unambiguous way to confirm that observed structures are veritable TNTs is to show that they are able to transfer material. By timelapse experiments we could demonstrate that TNT-like intercellular connections enable cytoplasmic continuity and transfer of cytoplasmic Dendra2 protein, suggesting that they should be open-ended and hence represent canonical TNTs. Indeed, Dendra2 protein molecular weight is 26 kDa which makes it impossible to pass through the gap junctions that allow transport of molecules smaller than 1 kDa. The fact that the speed of signal acquisition by the acceptor cell is almost equal to the speed of signal intensity loss from the donor cell, suggests direct cytoplasm transfer that can happen only in the case of open-ended connection. Moreover, we successfully demonstrated that TNTs in zebrafish facilitate the transfer of organelles, including mitochondria and early endosomes, which represents one of the most distinctive functions of TNTs.

Overall, our study represents the first demonstration of functional TNTs in the living zebrafish embryo. The identified structures share multiple morphological and dynamic characteristics with TNTs described in cell culture and enable the transfer of cellular material, which is to date the only reliable criteria to classify a given protrusion as a tunneling nanotube. Our results open the way to further investigations on the role of TNTs in the developing embryo. Of interest, several studies, including from our group ^34–37^, support potential implication of TNTs in various developmental processes, such as coordinating cell movement, cell differentiation, and cell proliferation. The current demonstration of the presence of these structures in living organisms strongly indicates that cells have the ability to communicate directly and selectively over considerable distances within tissues. TNT-mediated communication may encompass molecular, electrical, and mechanical signals and could potentially work alongside morphogen gradient signaling during embryonic development ^14^. Employing a targeted disruption of TNT formation in living organisms would be an effective strategy to assess the role of TNT-based communication. However, these studies will require better and more specific tools as well as further improved 3D imaging systems.

## Author contributions

Conceptualization, O.K., A.P., S.A., F.D.B. and C.Z.; Methodology, O.K., S.L., A.P. and S.A.; Investigation, O.K., S.L. and I.P.; Formal Analysis, O.K. and S.L.; Writing – Original Draft, O.K.; Writing – Review & Editing, O.K., S.L., A.P., S.A., F.D.B. and C.Z.; Supervision, C.Z.; Funding Acquisition, C.Z.

## Acknowledgements

Authors would like to acknowledge the help of the following Institut Pasteur facilities: Zebrafish Projects Hub, Photonic BioImaging platfrom, Biomaterials and Microfluidics platform, Image Analysis Hub, Bioinformatics and Biostatistics Hub. Authors thank Frédéric Rosa for his kind advice and help with the assembly of *pXT7-lyn-mKATE-E2A-phb-EGFP* plasmid. Authors would like to thank all members of the Zurzolo group for fruitful discussions and Reine Bouyssie, a member of the administrative staff of the Membrane Traffic and Pathogenesis Unit, for her continuous support. Authors would like to thank Janelia Research Campus Advanced Imaging Center members Teng-Leong Chew and Chad Hobson for valuable insights on the project.

This work was supported by grants Equipe FRM #EQU202103012692 and ANR-20-CE13-0032-01 to Chiara Zurzolo. Olga Korenkova was supported by the Pasteur-Paris University (PPU) International PhD Program and by FRM (Fondation Recherche Médicale) End of thesis grant #FDT202204014868. Shiyu Liu is supported by the “Programme Explore Cancer” de l’Institut Pasteur.

## Declaration of interests

The authors declare no competing interests.

## Methods

### Zebrafish husbandry

AB wild-type zebrafish (Danio rerio) were maintained in 3,5L tanks at a maximal density of seven per liter, in 28.5°C and pH 7.4 water, on a 14 h light/10 h dark cycle following to European Union guidelines for handling of laboratory animals. All procedures of animal caring were carried out in accordance with the official regulatory standards of the department of Paris and conformed to French and European ethical and animal welfare directives.

### Plasmids

The following plasmids were used to make mRNA for micro-injections in zebrafish embryos: *pXT7-lyn-mKATE-E2A-phb-EGFP*, *pDEST-LifeAct-mKATE-E2A-CEP55-EGFP*, *pDEST-LifeAct-mKATE-E2A-EGFP-tubulin*, *pDEST-lyn-Cerulean*, *pHTC-lyn-HaloTag*, *pDEST-LifeAct-mKATE-E2A-EGFP-EEA1*, *pDEST-Dendra2*, *pDEST-zEps8-HaloTag*, *pDEST-mEps8dCAP-HaloTag*.

All plasmids were assembled using NEBuilder® HiFi DNA Assembly Master Mix (New England Biolabs). *pXT7* and *pDEST* vectors were a kind gift from the group of Laure Bally-Cuif, *pG1-flk1-eGFP-tubulinα2* was a kind gift from Anne Schmidt, *pENTR_Mm_CEP55(WT)* was a gift from Guillermo de Cárcer & Marcos Malumbres (Addgene plasmid # 136333; http://n2t.net/addgene:136333; RRID:Addgene_136333), *actc1b-mKate2-EEA1* was a gift from Rob Parton (Addgene plasmid # 109508; http://n2t.net/addgene:109508; RRID:Addgene_109508), PhOTO-M (pMTB:memb-Dendra2-2A-H2B-Cerulean) was a gift from Periklis Pantazis (Addgene plasmid # 92401; http://n2t.net/addgene:92401; RRID:Addgene_92401), *pHTC HaloTag*® *CMV-neo* vector was purchased from Promega, *pXT7-lyn-mKATE-E2A-phb-EGFP* was kindly assembled by Frédéric Rosa.

Capped mRNA was prepared from purified linearized plasmids, using mMessage mMachine SP6 or T7 Transcription Kits (Invitrogen), following manufacturer’s protocol. Use of E2A sequence in the plasmids gives a possibility to inject single mRNA that will be transcribed into two separate proteins.

### Embryo micro-injections

For visualization of TNT-like connections, zebrafish embryos were micro-injected with mRNA at 16-cell stage at concentration of 100 ng/μl, into one of the central cells. For clonal expression, injection of *lyn-Cerulean* mRNA (100 ng/μl) at 8-cell stage was followed by injection of *lyn-mKATE-E2A-phb-EGFP* mRNA (100 ng/μl) at 16-cell stage. For Eps8 overexpression, embryos were injected at 1-cell stage with *Eps8-HaloTag* mRNA (100 ng/μl) and further injected with *lyn-mKATE-E2A-phb-EGFP* mRNA (50 ng/μl) at 16-cell stage. Embryos injected with only *lyn-mKATE-E2A-phb-EGFP* mRNA (100 ng/μl) at 16-cell stage were used as control.

### Embryo mounting and image acquisition

Imaging was performed live at gastrula stage of development (7-9 hpf) using Nikon Ti2E Spinning disk microscope with 40x NA 1.15 WD 0.6 water immersion objective. Prior to imaging, embryos were dechorionated at 5-6 hpf, by treatment with 0,5 mg/ml pronase at 25°C for 2 minutes, followed by manual removal of chorions in agarose-coated dishes using Dupont N5 forceps. Dechorionated embryos were mounted into custom-made agar chambers in a 35 mm glass bottom μ-dish (Ibidi) and oriented for the animal pole to face the coverslip.

Preparation of agar chambers: agar wells were prepared using a custom made PDMS (Polydimethylsiloxane) mold, designed according to Wühr et al. ^38^, with slight modifications. In short, a mold with 16 wells, with sides from 550 μm to 700 μm, was placed in a 35 mm glass bottom μ-dish. Wells were created by pouring 1,5 ml of 2% melted agarose into the μ-dish with the mold inside. After agar solidifying, the mold was carefully removed.

To visualize HaloTag expression, dechorionated embryos were incubated with HaloTag 647 ligand (Promega) at 28°C for 1 hour, followed by washing in E3 medium and mounting. For CK666 experiments, embryos were mounted into E3 medium containing either DMSO (control), or 50 μM CK666, or 200 μM CK666 and incubated for 1 hour at 28°C prior to imaging.

### Dendra2 photoconversion and time-lapse imaging

Cytoplasmic *dendra2* mRNA was co-injected together with *lyn-HaloTag* mRNA at 16-cell stage. Dechorionated embryos were incubated with HaloTag 647 ligand (Promega) at 28°C for 1 hour, followed by washing in E3 medium and mounting. Photoconversion of Dendra2 was performed using the FRAP module of Nikon Ti2E Spinning disk microscope, by 0,5 second exposure to 405 nm laser (at 25% laser power). ROI was imaged for 30 minutes with the time interval of 40 seconds between the stacks.

For all presented time-lapse recordings, time interval was kept in the range of 20-45 seconds between the stacks, representing the best time resolution available on the system.

### Image analysis and statistics

3D renderings of acquired Z-stacks were generated using the Imaris® software (version 9, Bitplane). The total number of labelled cells in the field of view was counted manually using Imaris Spots function. The length of every TNT-like connection was measured using Imaris Measurement points tool. The protrusion was annotated as a TNT-like connection if it was connecting two labelled cells, seemed continuous and had length above 5 μm. Every image represented separate embryo and had 50-100 labelled cells in the field of view, 20-40 images per group were analyzed.

For Dendra2 photoconversion experiment analysis, SUM intensity projection of the movie was used to measure mean fluorescent intensity (MFI) of donor cell, acceptor cell and 3 neighboring cells using Fiji software. Measured MFI was corrected for bleaching, using the bleaching speed calculated from another group of cells converted in the same embryo. For the visualization on the graph, corrected MFIs were normalized to the first time point MFI of the donor cell.

Statistical analysis was performed using Microsoft Excel and GraphPad Prism. All the statistical tests performed were two-tailed, and their significance level was set at 5%. To compare relative numbers of connected cells Mann Whitney test and Kruskal Wallis test were used, to compare average lengths of connections t-test and one-way ANOVA were used, to compare length distributions chi-square test was used.

## Supplementary files’ legends

**Supplementary figure 1.**
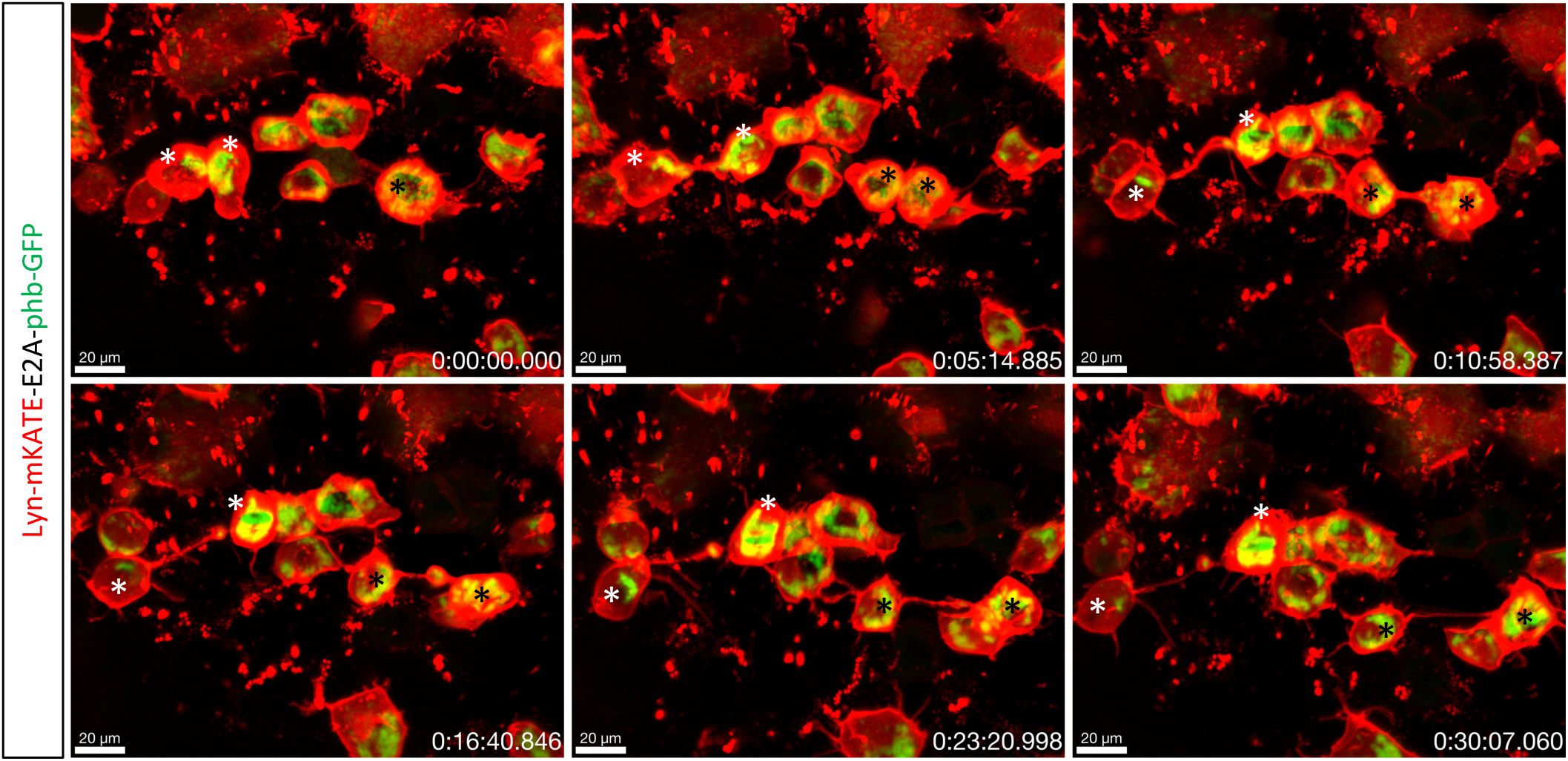
Stable cytokinetic bridges during zebrafish gastrulation. In several embryos we could observe unexpectedly long cytokinetic bridges. Timelapse recording at 8 hpf shows two pairs of daughter cells (white and black asterisks point corresponding pairs) connected by mitochondria-containing cytokinetic bridge that remains stable for at least 30 minutes of recording.

**Supplementary figure 2.**
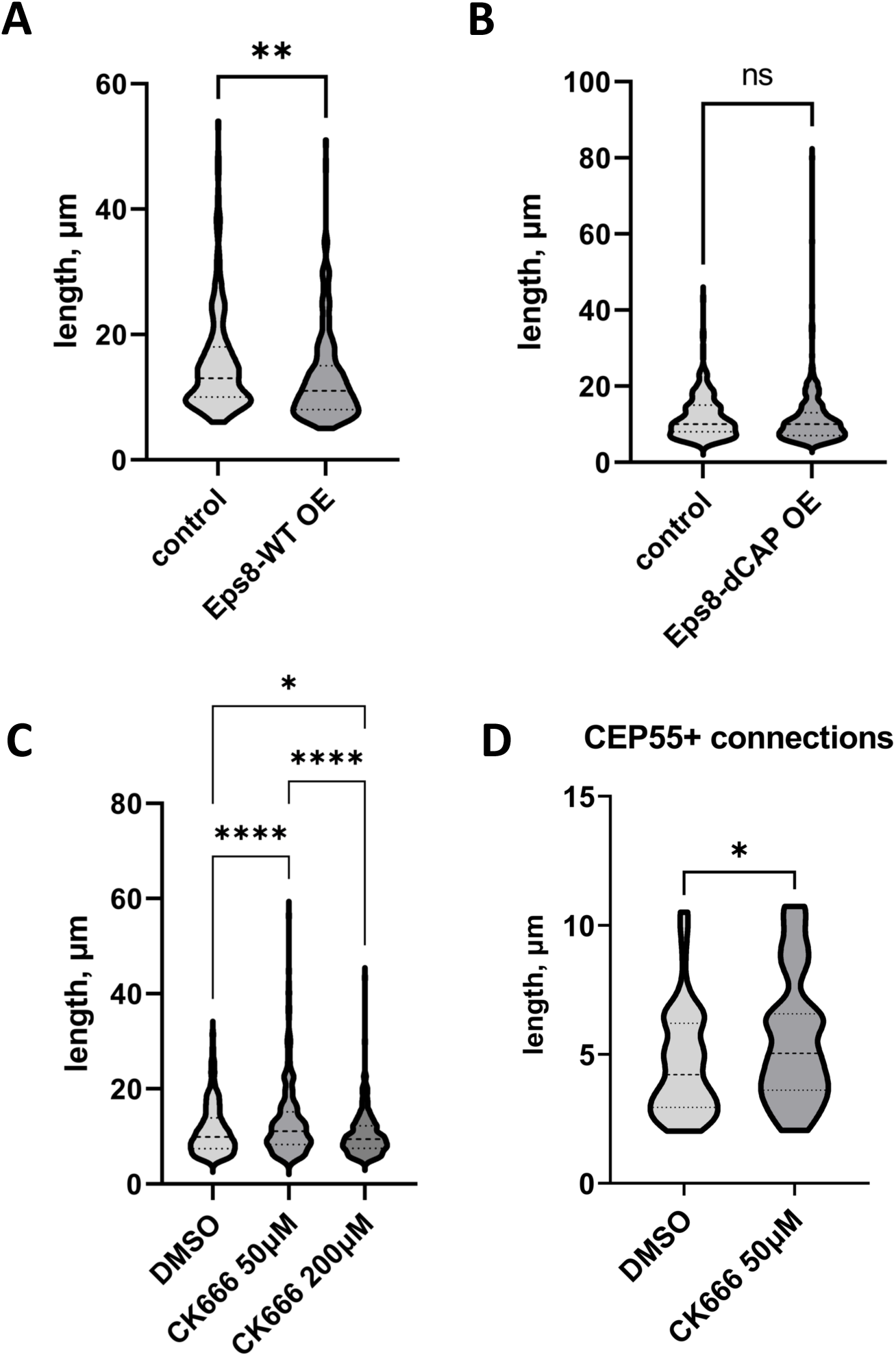
(A, B, C) Violin plots of length of TNT-like connections following treatments: Eps8-WT overexpression (A), Eps8-dCAP overexpression (B), CK666 treatment (C). (A) control group: n=220, mean 15.30 μm, Eps8-WT overexpression group: n=247, mean 12.99 μm. p= 0.0011, analysed by t-test. (B) control group: n=343, mean 11.77 μm, Eps8-dCAP overexpression group: n=489, mean 11.20 μm. p= 0.1717, analysed by t-test. (C) DMSO control group: n= 541, mean 11.29 μm, CK666 50 μM group: n=328, mean 12.90 μm, CK666 200 μM group: n=298, mean 10.34 μm. p-values are <0.0001 for DMSO vs. CK666 50μM, 0.0190 for DMSO vs. CK666 200μM and <0.0001 for CK666 50μM vs. CK666 200μM, analysed by one-way ANOVA. (D) Violin plot of length of CEP55-positive connections following CK666 treatment. DMSO control group: n= 82, mean 4.639 μm, CK666 50 μM group: n=51, mean 5.572 μm. p= 0.0192, analysed by t-test.

**Supplementary figure 3.**
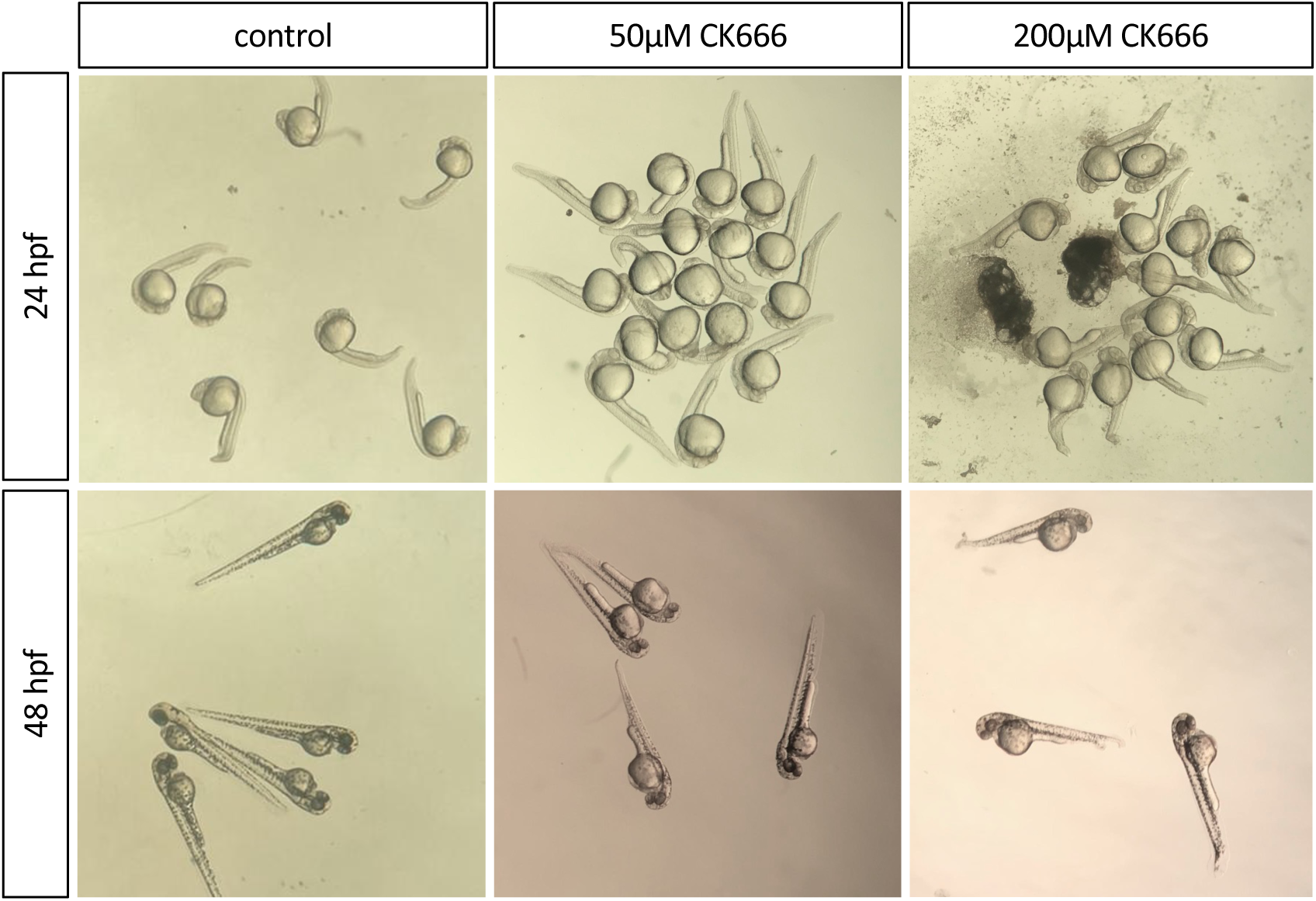
Toxicity of CK666 treatment on zebrafish embryos. Zebrafish embryos were treated with CK666 for 1 hour at the 6 hpf stage of development and left at 28°C for 24h and for 48h. Treatment with 50 μM CK666 did not alter the gross morphology of the embryos, while treatment with 200 μM CK666 resulted in embryonic death at early stages and twisting of the tail of the survived embryos at 48 hpf.

**Movie 1.** Stable cytokinetic bridges during zebrafish gastrulation.

Time-lapse recording of the zebrafish embryo at 8 hpf shows two pairs of daughter cells connected by mitochondria-containing cytokinetic bridges (yellow arrowheads) that remain stable for at least 30 minutes of recording.

**Movie 2.** TNT-like structures formed from two filopodia.

Time-lapse recording of the zebrafish embryo at 8hpf shows two neighboring cells extending filopodia (white arrowheads) that subsequently merge into single membranous connection that remains stable for at least 5 minutes.

**Movie 3.** TNT-like structures formed by cell dislodgement in parallel with cell division.

Time-lapse recording of the zebrafish embryo at 8hpf shows three neighboring cells forming TNT-like connections by cell dislodgement (yellow arrowheads), while the top cell is dividing (white asterisks). Daughter cells maintain connections with lower cells for at least 15 minutes (yellow arrowheads), while cytokinetic bridge between daughter cells is also visible (white arrowhead).

**Movie 4.** TNT-like connections elongate along each other.

Time-lapse recording of the zebrafish embryo at 8hpf shows two cells belonging to different cell populations (top red cell and bottom green cell), extending protrusions of corresponding colors (red and green arrowheads) that elongate along each other.

**Movie 5.** Zebrafish TNTs allow transfer of cytoplasmic Dendra2 protein.

Time-lapse recording of the zebrafish embryo at 8hpf shows two cells connected by a membranous TNT-like connection (yellow arrowhead). Cytoplasmic Dendra2 was photoconverted to red in the cell on the left (donor), recording shows that cell on the right (acceptor) receives red labeling within 20 minutes. In the end of the recording connection breaks (white arrowhead).

**Movie 6.** Zebrafish TNTs allow transfer of mitochondria.

Time-lapse recording of the zebrafish embryo at 8hpf shows two cells connected by a membranous TNT-like connection, which enables the transfer of mitochondria (white arrowhead).

**Movie 7.** Zebrafish TNTs allow transfer of early endosomes.

Timelapse recording of the zebrafish embryo at 8hpf shows actin-rich connection between two cells (white arrowhead) that allows the transfer of early endosomes (yellow arrowhead).

## Notes

### Competing Interest Statement

The authors have declared no competing interest.

